# Design of an Epitope-Based Peptide Vaccine against the Severe Acute Respiratory Syndrome Coronavirus-2 (SARS-CoV-2): A Vaccine-informatics Approach

**DOI:** 10.1101/2020.05.03.074930

**Authors:** Aftab Alam, Arbaaz Khan, Nikhat Imam, Mohd Faizan Siddiqui, Mohd Waseem, Md. Zubbair Malik, Romana Ishrat

**Author notes:** **Correspondence: Dr. Romana Ishrat, Ph.D.**, Centre for Interdisciplinary Research in Basic Science, Jamia Millia Islamia University, New Delhi-110025, India.

## Abstract

The recurrent and recent global outbreak of SARS-CoV-2 has turned into a global concern which has infected more than 19-million people all over the globe, and this number is increasing in hours. Unfortunate no vaccine or specific treatment is available, which make it more deadly. A vaccine-informatics approach has shown significant breakthrough in peptide-based epitope mapping and opens the new horizon in vaccine development. In this study, we have identified a total of 15 antigenic peptides (including T and B cells) in the surface glycoprotein of SARS-CoV-2 which showed non-toxic nature, non-allergenic, highly antigenic and non-mutated in other SARS-CoV-2 virus strains. The population coverage analysis has found that CD4^+^ T-cell peptides showed higher cumulative population coverage over to CD8^+^ peptides in the 16 different geographical regions of the world. We identified twelve peptides *(LTDEMIAQY, WTAGAAAYY, WMESEFRVY, IRASANLAA, FGAISSVLN, VKQLSSNFG, FAMQMAYRF, FGAGAALQI, YGFQPTNGVGYQ, LPDPSKPSKR, QTQTNSPRRARS and VITPGTNTSN*) that are 80% - 90% identical with experimentally determined epitopes of SARS-CoV, and this will likely be beneficial for a quick progression of the vaccine design. Moreover, docking analysis suggested that identified peptides are tightly bound in the groove of HLA molecules which can induce the T-cell response. Overall this study allows us to determine potent peptide antigen targets in surface glycoprotein on intuitive grounds which open up a new horizon in COVID-19 research. However, this study needs experimental validation by in vitro and in vivo.

## Introduction

History is witness to the fact that whenever a major disease comes, it brings tremendous destruction with itself, and these destructions are physical and economical. Previously, Cholera, Smallpox, Bubonic plague and Influenza are some of the most brutal killers in human history and killed millions of people all over the world. Nowadays, the world is scared of Coronavirus disease (COVID-19), and its outbreak continues to spread from China to all over the world, and we do not yet know when it will stop. It is a contagious disease caused by SARS family virus named ‘Severe acute respiratory syndrome coronavirus 2 (SARS-CoV-2)’, which has not been previously identified in humans. The initial symptoms of COVID-19 are dry cough, throat infection, fever, breath shortness, loss of taste or smell, rashes in skin, conjunctivitis, tiredness, muscles pain, diarrhoea; most of the individuals (about 80%) recover without any special treatment, but older people and a person with pre-existing medical conditions or comorbidities (cardiovascular disease, diabetes, lungs disease, and cancer) are more likely to develop serious illness. The incubation period for the infection with SARS-CoV-2 ranges from 2 to 14 days after its exposure[1]. The transmissibility of the virus is shown by its reproductive number (R0), and it varies area to area. The World Health Organization (WHO) has estimated R0 between 1.4 and 2.5, but some other studies have estimated 2.24 to 3.58 of COVID-19[2,3].

In just nine months, the COVID-19 that originally emerged from the Wuhan province of China poses a major public health and governance challenges. The cases have now spread in 213 countries and territories around the globe. Till 24^th^ October 2020, more than 42.5 million infected individuals with over 1,134,940 deaths around the globe have been reported to WHO (WHO COVID-19 Dashboard), and these numbers are rapidly increasing in hours. At present, no vaccine or specific treatment is available; However, the World Health Organization listed (as per 19 Oct 2020) more than 200 vaccines in development at various stages (Pre-clinical evaluation-154, clinical evaluation-44). The vaccine candidates which are listed in the clinical evaluation stage seven have reached phase III trail, includings Ad5-nCoV (CanSino Biologics, CHINA), AZD1222 (The University of Oxford; AstraZeneca; IQVIA; Serum Institute of India), CoronaVac (Sinovac), JNJ-78436735 or Ad26.COV2-S (Johnson & Johnson), mRNA-1273 (Moderna), Unknown vaccine (No name announced by Wuhan Institute of Biological Products; China National Pharmaceutical Group (Sinopharm) and NVX-CoV2373 (Novavax). Further, The University of Melbourne and Murdoch Children’s Research Institute; Radboud University Medical Center; Faustman Lab at Massachusetts General Hospital’s BCG live-attenuated vaccine also in Phase II/III combined phase. The AstraZeneca/ University of Oxford vaccine candidate (AZD1222) looks the most promising vaccine candidates, which is in Phase II/III combined phase. (WHO: Draft landscape of COVID-19 candidate vaccines).

Moreover, there are no chemotherapeutic agents available to curb this menace; however, few agents are being used including natural compounds[4–6], western medicines[7,8] and traditional Chinese medicines (TCN)[9,10] that may have potential efficacy against the SARS-CoV-2. Moreover, other drugs like interferon α (IFN-α), lopinavir/ritonavir, chloroquine phosphate, ribavirin, favipiravir, disulfiram, arbidol and hydroxychloroquine are recommended as the tentative treatments of COVID-19 [11,12]. Currently, there is no specific treatment/medicine or vaccine to cure COVID-19, so there is an urgent need to develop new vaccines or drugs against this deadly disease. In this way, integration of computational techniques provides a novel approach to integrating vaccine-informatics approach for the development of vaccines. These methods had earlier been used in the development of vaccines against the several diseases including Dengue[13], Malaria[14], Influenza[15], Multiple sclerosis[16], and tumour[17]. However, this approach generally works through the identification of MHC-1 & II molecules and T-cell epitopes (CD8^+^ & CD4^+^)[18], which particularize the selection of the potential vaccine agents related to the transporter of Antigen Presentation (TAP) molecules[19,20].

The beginning of 2020 has seen the emergence of the deadly COVID-19 pandemic, and currently, we are drowning with the enormous amount of articles with their probable epitope-based peptide vaccine for COVID-19 in which the B and T-cell epitopes have been analyzed and anticipated the possibility of antigenic epitopes which can be used to design a novel vaccine candidate against the SARS-CoV-2[21–23]. In the current study, we have also predicted epitope-based vaccine candidates against the SARS-CoV-2 using the systematic vaccine-informatics approach. We considered surface glycoprotein of SARS-CoV-2 due to its specific functions; SARS-CoV-2 uses surface spike protein to mediate entry into host cells. To fulfil its purpose, SARS-CoV-2 spike binds to receptor ‘hACE2’ through its receptor-binding domain (RBD) and is proteolytically triggered by human proteases[24,25]. Notably, we not only prognosticate the most potential vaccine candidate but also crosschecked the resemblance, congruity and the compatibility of these selected epitopes with human to circumvent any possible risk of autoimmunity. Additionally, we had checked the resemblance of our epitopes with those which are already experimentally verified epitopes of different organisms including SARS-CoV, which not only makes our study more precise and noteworthy but also expands our views for vaccine-informatics approach in planning for the next global pandemic. Our work can save the time needed to screen a large number of possible epitopes compared with experimental techniques and also guide the experimental work with high confidence of finding the desired results in vaccine development.

## Materials and methods

### 1. Sequence Retrieval and Analysis

The surface glycoprotein (SG) sequence (ID:-QHO62112.1) was obtained from the NCBI gene bank database (https://www.ncbi.nlm.nih.gov/gene/) in the FASTA format. Additionally, we check the sequence similarity of peptide sequences with other SG proteins of other SARS-CoV-19 isolates from different geographical regions using Clustal Omega tool[26] to analyze the variation in epitopes-sequences that can determine us whether the epitopes are conserved or have altered peptide ligands.

### 2. T-Cell Peptides (Epitopes) Prediction

The NETCTL_1.2 server[27] was used to identify the CD8^+^ T-Cell peptides at set threshold value 0.95 to sustain the sensitivity and specificity of 0.90 and 0.95, respectively. We used all the expanded 12 MHC-I supertypes (including *A1, A2, A3, A24, A26, B7, B8, B27, B39, B44, B58 and B62)* and incorporated the peptide prediction of *MHC class - I* binding molecules and Proteasomal C-terminal cleavage with TAP transports efficiency. These results were accomplished by weighted TAP transport efficiency matrix. Then MHC-I binding and proteasomal cleavage efficiency were used to obtain the total scores and translated it into the sensitivity and specificity. We selected peptides as the epitope candidate on the basis of the overall score. Further, we check the peptides binding to *MHC class - I* molecules by using MHC-I binding predictions tool[28]. The predicted output was given in units of IC_50_ *nM* (IC_50_ values < 50 *nM* = high affinity, IC_50_< 500 *nM* moderate affinity and IC_50_ < 5000 *nM* low affinity)[29]. The MHC-NP tool used for assesses the probability of naturally processing and binding of peptides to the given MHC molecules. This tool predicts naturally processed epitopes based on physiochemical properties and comparison of residual position with experimentally verified epitopes[30]. Further, we identified CD4^+^ T-Cell peptides with IC50 ≤ 100 ((IC_50_ values < 50 *nM* = high affinity, IC_50_< 500 *nM* moderate affinity and IC_50_ < 5000 *nM* low affinity) by using MHC-II binding predictions tool[31].

### 3. B-Cell Peptides (Epitopes) Prediction

The identification of B-cell peptides (Epitopes) in the surface glycoprotein of SARS-CoV-2 was accomplished to find the potent antigen which gives an assurance of humoral immunity. Here, we used Antibody Epitope Prediction tool[32] to find the B-cell antigenicity with classical propensity scale methods including *Bepipred Linear Epitope Prediction 2.0* [33], *Chou & Fasman Beta-Turn Predictio*n[34], *Emini Surface Accessibility Prediction*[35], *Karplus & Schulz Flexibility Prediction*[36], *Kolaskar & Tongaonkar Antigenicity*[37] and *Parker Hydrophilicity Prediction*[38]. In this study, we selected that region in the protein sequence, which was cross-referenced, and common findings were considered as the B-cell antigenic region based on the above six classical propensity scale methods.

### 4. Epitope Conservancy and Immunogenicity Analysis

The epitope-conservation defined the degree of a resemblance betwixt the peptide and query sequences. Hence, we used the Epitope Conservancy Analysis (ECA) tool [39] to find conservancy of our peptides among the other SARS coronaviruses, Bat SARS-like coronaviruses and Bat coronaviruses. Additionally, the immunogenicity prediction can reveal the degree of efficiency for individual peptides to produce an immunogenic response. In our study, we used Class I Immunogenicity[40] tool to predict the immunogenicity of the MHC class-I binding peptides.

### 5. Peptide Structural Modelling and Retrieval of Human Leukocyte Antigen (HLA) Molecules

The structure of CD^+^8 and CD^+^4 T-cell peptides were modelled by a biased modeling method using an online server PEP-FOLD 3.5 server at RPBS MOBYL portal [41]. The crystal structure of SARS-CoV-2 spike glycoprotein (6VSB-A) were used as the reference model together with a mask representing the structure fragments for which the local conformation of this structure has been imposed. Additionally, The structure of HLA molecules including HLA-B*53:01 (PDB ID: *1A1M*)[43], HLA-B*44:03(PDB ID: *4JQX*)[44] and HLA-DRB1*01:01(1AQD)[45] were retrieved from the Protein Data Bank (PDB)[46]. The structure of peptides and HLA molecules are depicted in Figure 1.

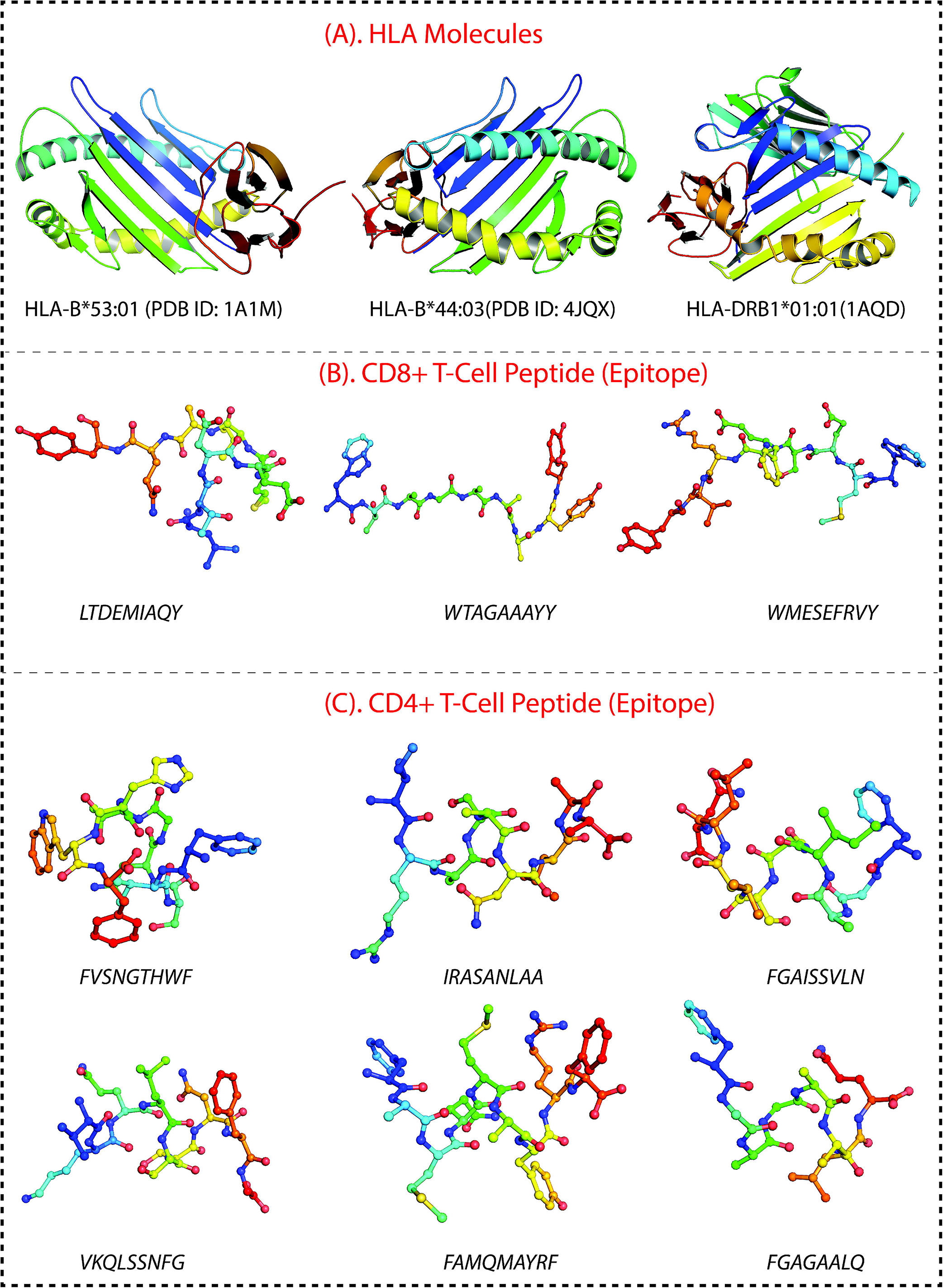

### 6. Population Coverage Analysis (PCA)

The Population Coverage Analysis tool[47] gives the idea about the probable response of each peptide in different countries of the world based on peptide-HLA data genotypic frequency. Our CD8^+^ T-cell (three peptides) and CD4^+^ T-cell (six peptides) peptides and their respective HLA alleles were used for PCA. We selected 16 geographical areas (115 countries and 21 different ethnicities grouped.) were *East Asia, Northeast Asia, South Asia, Southeast Asia, Southwest Asia, Europe, East Africa, West Africa, Central Africa, North Africa, South Africa, West Indies, North America, Central America, South America and Oceania*. These 16 geographical regions cover the HLA allele frequencies and associated data for different individual populations from the most popular countries.

### 7. Predicted Peptides vs. Epitope Database and Peptide Screening for Autoimmune, Allergic and Toxic Response

In this section, we used predicted peptides (including T and B–cell peptides) to search for homologous epitopes at 80% to 90% identity in Epitope database[48,49]. The current limited accessible data isn’t sufficient to recognize antigenic region in spike proteins of SARS-CoV-2 by human immune responses. Herein homologous epitopes (derived from other pathogens) would expedite the evaluation of vaccine candidate immunogenicity and would aid in vaccine designing. Furthermore, to surveil the prospective consequences of mutational events and epitope escape as the virus is transmitted through human populations[50]. Based on initially full-length genomic phylogenetic analysis, the precursory studies suggested that SARS-CoV-2 is quite similar to SARS-CoV, and the putatively same cell entry mechanism and human cell receptor usage. And due to conspicuous resemblance amid these viruses, the previous study that gave us a basic understanding of protective immune responses against SARS-CoV may potentially be leveraged to aid vaccine development against the SARS-CoV-2[51]. This study also helps to check peptides identity with human proteome because there is a chance of an autoimmune response due to any kind of molecular mimicry between the predicted peptides (epitopes) and the human proteome. Moreover, we checked the allergic and toxic nature of predicted peptides using AlgPred[52] and ToxinPred[53] tools, respectively.

### 8. Molecular Docking Studies

In this study, molecular docking studies produced important information regarding the orientation-pattern of the peptides in the binding groove of the HLA-molecules as well as which residues are actively involved in the binding. In this study, we selected three CD8^+^ T-cell peptides (*LTDEMIAQY, WTAGAAAYY* and *WMESEFRVY)* and five CD4^+^ T-cell peptides (*IRASANLAA, FGAISSVLN, VKQLSSNFG, FAMQMAYRF* and *FGAGAALQ*) for molecular docking against the HLA molecule including HLA-B*53:01 (PDB ID: *1A1M*), HLA-B*44:03 (PDB ID: *4JQX*) and HLA-DRB1*01:01 (PDB ID: 1AQD). We used glide module[54–56] of Schrödinger suite for peptides-HLA molecules docking. All peptides were prepared using LigPrep module of Schrodinger suite and docked in binding site of protein using SP-peptide mode of Glide. Receptor grid was generated using Receptor grid generation in the Glide application by specifying the binding (active) site residues, which was identified by SiteMap tool[57]. The docked conformers were evaluated using Glide (G) Score and the best docked pose with lowest Glide score value was recorded for each peptide.

## Results

### 1. Sequence Retrieval and Structure Prediction

Viral glycoproteins have a major role in its pathogenesis. The main goal of viral infection is to recognise and bind a receptor on the cell surface of the host. It is considered as the surface glycoprotein play an important role in immunity and infection. We retrieved the envelope surface glycoprotein of SARS-CoV-19 from the NCBI gene bank database (ID: QHO62112.1). Moreover, the sequence similarity of query proteins and peptides were done with the other SG proteins of other SARS-CoV-19 isolates from various regions of the world (Including China, Columbia, Japan, Malaysia, Israel, Iran, India, Sri Lanka, Vietnam, South Korea, Pakistan, USA, Hong Kong, Taiwan, Spain, South Africa, Serbia, Greece, Nederland, France and the Czech Republic) and found that all the predicted peptides are conserved in all of the isolates (As per 10^th^ August 2020)<u>(Supplementary data-1).</u>

### 2. Identification of T-cell Epitopes from SG protein of SARS-CoV-19

The NETCTL server predicted several peptides in SARS-CoV-19 surface glycoprotein but only nine most potent peptides were chosen which have a high combinatorial score. We considered only those alleles of MHC-I class for which the peptides showed higher binding affinity (*IC50 ≤ 400nm*). Proteasomes played a key role in cleaving the peptide bonds and converting the protein into a peptide. The total score of each peptide–HLA interaction was taken into consideration, and a greater score meant greater processing efficiency. The three peptides *LTDEMIAQY* (P1), *WTAGAAAYY* (P2) and *WMESEFRVY* (P3) among nine were found to bind with most of the MHC Class-I molecules, including *HLA-B*15:01*, *HLA-B*53:01*, *HLA-A*68:02*, *HLA-B*44:03* and *HLA-B*57:01* but peptides P1, P2 and P3 had a maximum probable value of 0.8203, 0.8185 and 0.7539 for the *HLA-B*53:01*, *HLA-B*44:03* and *HLA-B*44:03* respectively. These peptides (P1, P2 and P3) also have a maximum identity (100%) for the conservancy. Moreover, we made the immunogenicity prediction of peptides and got the highest pMHC-I immunogenicity scores of 0.02757 (P1), 0.15259 (P2) and 0.14153 (P3). The details are given in Table 1. Additionally, we identified 162 CD4^+^ T-Cell peptides (Epitopes) with *IC50 ≤ 100*; however, only six peptides (*FVSNGTHWF, IRASANLAA, FGAISSVLN, VKQLSSNFG, FAMQMAYRF,* and *FGAGAALQ*) were found to interact with most of the HLA-DRB-1 molecules, the details are given in Table 2.

### 3. Identification of Linear B-cell Epitopes from SG protein of SARS-CoV-19

The B-cell epitopes comprise of peptides which can easily be used to take the place of antigens for immunizations and antibody production. In this study, we used an amino acid scale-based method in B-cell antibody epitope prediction tool in which we predict linear epitopes from protein sequence. We found six B-cell linear epitopes (including *LTPGDSSSGWTAG, YQAGSTPCNGV, YGFQPTNGVGYQ, VITPGTNTSN, QTQTNSPRRARS* and *LPDPSKPSKR*) in the surface glycoprotein of SARS-Cov-19 which may be capable of inducing the desired immune response as B-cell epitopes (Figure 2).

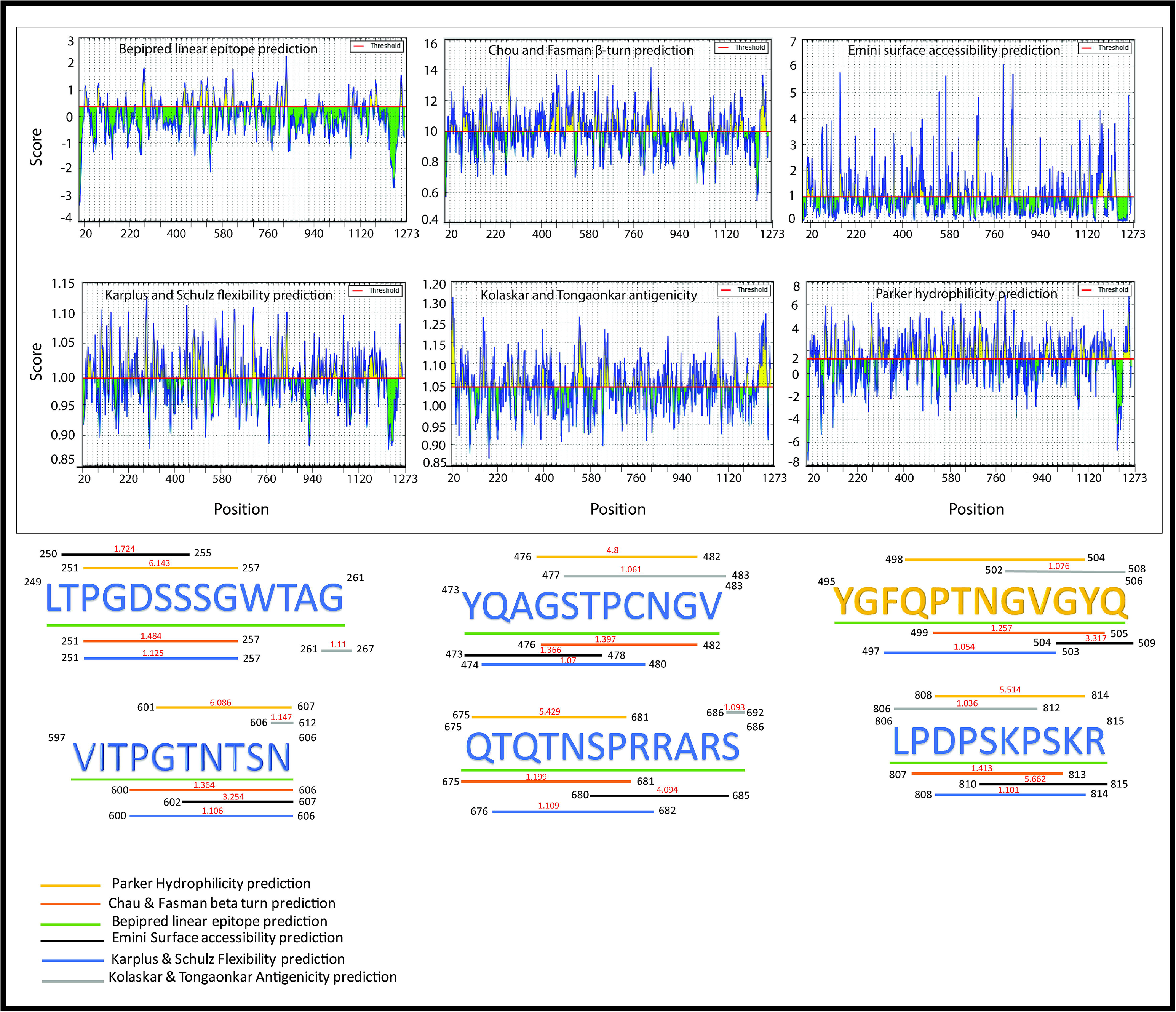

### 4. Population Coverage Analysis (PCA)

The population coverage analysis calculates the expected response of predicted peptides (including B and T-cells) in populations from different geographical areas. In this study, PCA for 16 different geographical regions was carried out for the predicted peptides by considering all the MHC class-I & II molecules. These results suggested that the expected response of these peptides is varying for populations residing in different geographical regions, shown in Figure 3. The tool predicted average population coverage of 21.50% and 51.09% for MHC Class-I & II binding peptide fragments respectively. The PCA of MHC class II binding peptide fragments of SARS-CoV-19 surface glycoprotein revealed maximum population coverage, e.g., 75.57%, 73.61% and 72.91% for North America, Europe and East Asia populations.

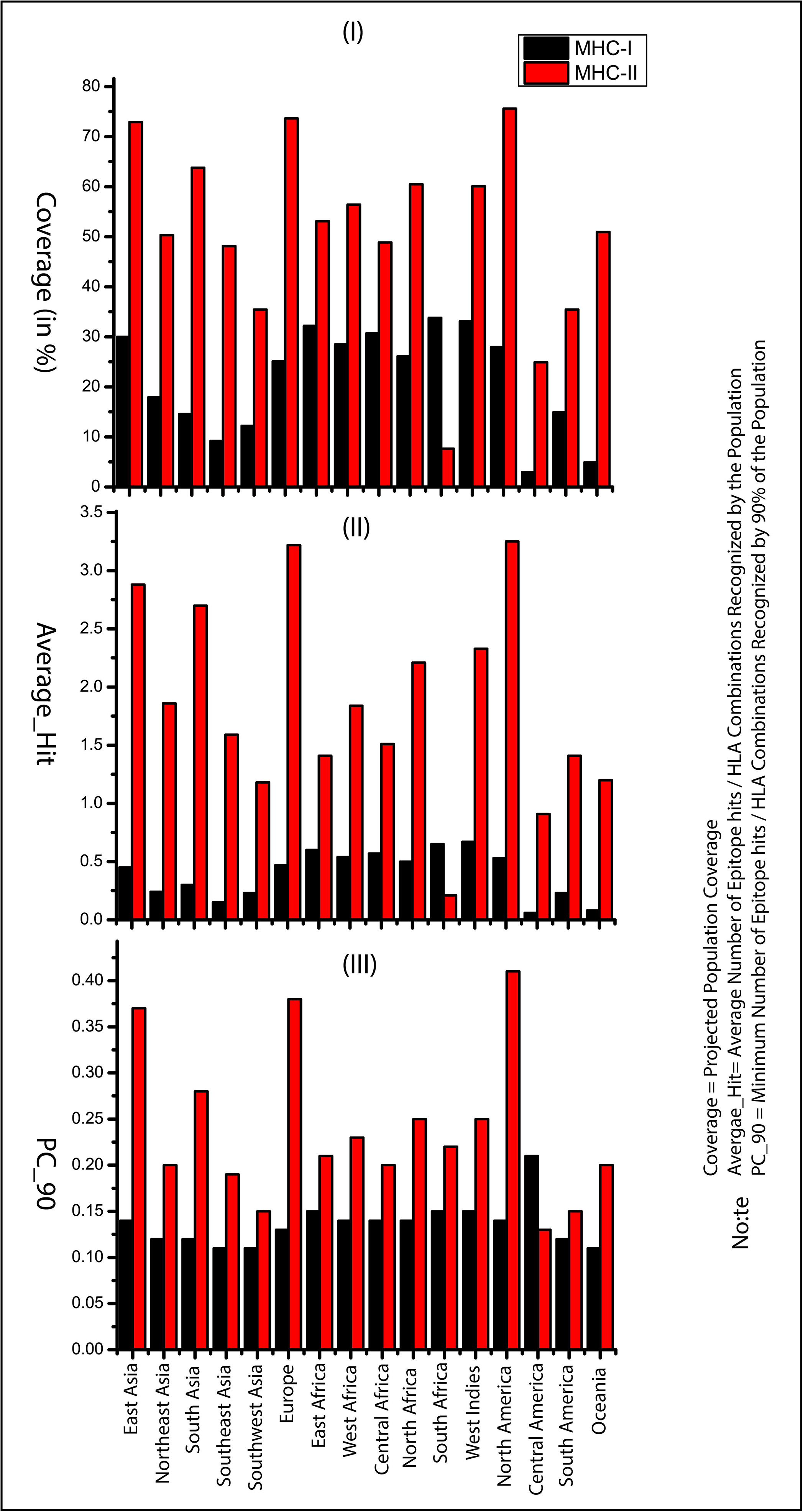

### 5. Autoimmune, Allergic and Toxic Response

In the past, there are many cases have been reported for the development of autoimmune diseases like systemic lupus anaemia, mysthemia gravis (hepatitis B), multiple sclerosis (swine flu), diabetes mellitus (Mumps) and Gullian Barr syndrome (Influenza and oral polio vaccine)[58,59]. To avoid autoimmunity, it becomes important to discard those peptide agents which are similar to the human proteome. So, we have mapped all the predicted peptides sequences against the human and other viruses, including SARS-CoV, which has a maximum sequence identity to SARS-CoV-2 and is the best-characterized coronavirus in respect to epitopic responses. We have identified most of the peptide sequences (including B and T-cell peptides) were found to similar with experimentally determined epitopes of SARS-CoV virus except two peptides (*FVSNGTHWF and LTPGDSSSGWTAG*) which resemble with Mus *musculus* and human’s proteome; these peptides were eliminated from further study due to autoimmunity risk. The details of identified peptides and their resemblance with another organism’s proteome are given in Table 3. Next, we have screened all the peptides to check their allergic and toxic nature, and all these peptides were found non-Allergen as well as non-toxic in nature

### Peptide-HLA Interaction Analysis

To ensure the interaction between the CD^+^8 T-cell peptides (*LTDEMIAQY*, *WTAGAAAYY* and *WMESEFRVY*) and HLA molecules (HLA-B*53:01(1A1M), HLA-B*44:03(4JQX) and HLA-B*44:03(4JQX) respectively), we performed molecular docking analysis and found that the peptide *LTDEMIAQY* bind with HLA-B*53:01 having good docking score −9.54 kcal/Mol. Similarly, *WTAGAAAYY* and *WMESEFRVY* bind with HLA-B*44:03 having binding affinity −8.80 kcal/Mol and −9.22 kcal/Mol, respectively. Moreover, all the CD^+^4 T-cell peptides were binds into the groove of HLA-DR molecules (HLA-DRB1*01:01(1AQD)) with good docking score, e.g., −10.63 kcal/Mol (with *IRASANLAA*), −12.19 kcal/Mol (with *FGAISSVLN*), −8.74 kcal/Mol (with *VKQLSSNFG*), −8.59 kcal/Mol (with *FAMQMAYRF*) and −9.28 kcal/ Mol (with *FGAGAALQ*).

## Discussion

The development of vaccines refers to one of the most effective and cost-effective medical and public health achievements of all time. It is a very lengthy, complex, and costly process and requires the involvement of public and private combination. In each year, vaccination programs save over three million lives. The peptide as a choice of vaccine agent has made the astonishing move towards vaccine design against the viruses, bacteria and cancer. The peptide vaccine is often synthetic and mimics naturally occurring proteins from pathogens, and these peptides based vaccine have shown promising successful results in the past for disease like malaria, dengue, multiple sclerosis, Influenza. Besides these diseases, the peptide-based vaccines have also been developed against several types of cancer like colorectal cancer, myeloid leukaemia and gastric cancer[60–62]. The identification and design of immunogenic peptides (epitopes) is expensive as well as time-consuming. So vaccine-informatics approach has made it easy to identify potent peptides. In the present study, we have identified CD8^+^ T-cell (three peptides), CD4^+^ T-cell (six peptides) and B-cell linear peptides (six peptides) from SARS-CoV-19 surface glycoprotein; however, we were more emphasized to study T-cell peptides because vaccines against T-cell epitopes are more promising as they evoke a long-lasting immune response, with antigenic drift, an antigen can easily escape the memory response of antibody[63,64].

For the MHC class-I binding peptides, the immune response for the top five different geographical regions (highest population coverage range 30-34%) including South Africa_33.78% (South Africa), West Indies_33.12% (Cuba, Jamaica, Martinique, Trinidad and Tobago), East Africa_32.25% (Kenya, Uganda, Zambia and Zimbabwe), Central Africa_30.70% (Cameroon, Central African Republic, Congo, Equatorial Guinea, Gabon, Rwanda and Sao Tome & Principe) and East Asia_30% (Japan, South Korea and Mongolia). Similarly, for the MHC class-II binding peptides, the excepted immune response was found to be remarkable (PCA range 60%–76%) for North America (Canada, Mexico and the United States), Europe (Austria, Belarus, Belgium, Bulgaria, Croatia, Czech Republic, Denmark, England, France, Georgia, Germany, Greece, Ireland Northern, Ireland South, Italy, Macedonia, Netherlands, Norway, Poland, Portugal, Romania, Russia, Scotland, Serbia, Slovakia, Spain, Sweden, Switzerland, Turkey, Ukraine, United Kingdom and Wales), East Asia (Japan, South Korea and Mongolia), South Asia (India, Pakistan and Sri Lanka), North Africa (Algeria, Ethiopia, Mali, Morocco, Sudan and Tunisia) and West Indies (Cuba, Jamaica, Martinique, Trinidad and Tobago). Thus, our results suggested that MHC class-II binding peptides may be potent vaccine agents for different populations of the globe.

We know that vaccination is the utmost prevention of epidemiologic infectious diseases, but it has a low incidence of serious systemic adverse effects (like autoimmune diseases). We have predicted a total of 15 peptides (B and T-cell), but twelve peptides are very important including *LTDEMIAQY, WTAGAAAYY, WMESEFRVY, IRASANLAA, FGAISSVLN, VKQLSSNFG, FAMQMAYRF, FGAGAALQI, YGFQPTNGVGYQ, LPDPSKPSKR, QTQTNSPRRARS and VITPGTNTSN* because their peptide fragments are matching with experimentally identified epitopes of the glycoprotein of Severe acute respiratory syndrome-related coronavirus (<u>SARS-CoV</u>)[65–69], thus, these peptides are seemingly a more rational set of potential vaccine agents against the SARS-CoV-2.

Moreover, the other two peptides *LTPGDSSSGWTAG and FVSNGTHWF,* which shown resembles with Homo sapiens (human) and Mus musculus proteome, were eliminated from the study to avoid autoimmunity risk. Besides, we didn’t find any resemblance of one peptide (*YQAGSTPCNG)* with any organism’s proteome. Hence, this unique peptide can also be proposed as candidates for further studies in the area of vaccines development, therapeutic antibodies and diagnostic tools against the SARS-CoV-2.

As we know that HLA-alleles are polymorphic in nature; so peptide can interact with it. In our study, MHC class-I binding peptides (8-9 residues long) are binds in the wide groove (~1.25À) of HLA molecules (HLA-B*53:01 and HLA-B*44:03). Peptide (*LTDEMIAQY)* bind with HLA-B*53:01 by *ILE-66, THR-69, ASN-70, THR-73, GLU-76, ASN-77, TYR-84, ARG-97, TYR-99, TYR-123, THR-143, TRP-147, TYR-159* residues and other surrounding residues which gives it polymorphic nature lie at the interface of this helix and the bottom of the peptide-binding site. In docking with HLA-B*44:03, the peptides *WTAGAAAYY (*interacted with *GLU-76, THR-80, ALA-81, ILE-95, ASP-116,TYR-123, LYS-146, TRP-147, ALA-158, LEU-163 and GLU-166)* and W*MESEFRVY (*interacted with *THR-73 GLU-76 ARG-83, TYR-84, ARG-97, ASP-114, ASP-116, LYS-146 TRP-147, ALA-149, ALA-150 and LEU-156)* got expanded conformation within the same trench, and both peptides were surrounded by polymorphic region. Further, we check the interaction of MHC class-II binding peptides with HLA-DR molecules. All the five peptides are buried by interactions with the most important pocket residues.e.g. *GLN-09, PHE-13, ASN-62, VAL-65, GLN-70, ARG-71, TYR-78 etc.* and are preferred bind in the groove of HLA-DR molecules. The docking studies suggested that these peptides are preferably bound into the groove of respective alleles and can induce the CD^+^8 and CD^+^4 T-cell immunity.

In our study, integrated computational approaches have identified a total of 15 peptides (T and B cells) from SARS-CoV-19 surface glycoprotein, of which twelve peptides have resemblance with experimentally identified epitopes of SARS-CoV and other pathogens. Since there is currently no vaccine or specific treatment for COVID-19 and vaccine development is a long way from being translated into practical applications. So at this crunch situation, we have suggested that the predicted peptides (epitopes) would be capable to prompt an effective immune response as a peptide vaccine against the SARS-CoV-2. Consequently, these peptides may be used for synthetic vaccine design to combat emerging COVID-19. However, in vivo and in vitro, experimental validation is needed for their efficient use as a vaccine candidate.

## Author Contributions

RI and AA conceived the study design instructed on data analysis. AK and AA curated data and performed statistical analyses. N.I and MFS curated data and draw figures. M.W and MZM performed molecular docking studies. AA improved and revised the manuscript. All the authors read, edited and approved of the manuscript.

## Funding

This work did not receive any specific grant from funding agencies in public, commercial or not-for-profit sectors.

## Conflict of Interest

The authors declare that the research was conducted in the absence of any commercial or financial relationships that could be construed as a potential conflict of interest.

## Acknowledgements

The R.I and A.K are grateful to the Centre for Interdisciplinary Research in Basic Sciences (CIRBSc), Jamia Millia Islamia-110025 for providing the research infrastructure. All the authors also grateful for School of Computational and Integrative Sciences (SC&IS), JNU, New Delhi for providing Glide, Schrödinger software. N.Imam is thankful to the Indian Council of Medical Research (ICMR), New Delhi, India. Md.F.Siddiqui is grateful to Prof. Muratov Zhanybek Kudaibakovich and Dr. Syed Ali Abbas from Osh State University, Kyrgyzstan for their guidance and support. Aftab Alam acknowledges the Department of Health Research (DHR) New Delhi, India for the award of “Young Scientist” fellowship (R.12014/06/2019-HR).

## Notes

### Competing Interest Statement

The authors have declared no competing interest.

## References

1. Guo Y-R, Cao Q-D, Hong Z-S, et al. The origin, transmission and clinical therapies on coronavirus disease 2019 (COVID-19) outbreak - an update on the status. Mil. Med. Res. 2020; 7:11

2. Zhao S, Lin Q, Ran J, et al. Preliminary estimation of the basic reproduction number of novel coronavirus (2019-nCoV) in China, from 2019 to 2020: A data-driven analysis in the early phase of the outbreak. Int. J. Infect. Dis. 2020; 92:214–217

3. Alam A, Siddiqui MF, Imam N, et al. Covid-19: current knowledge, disease potential, prevention and clinical advances. Turk. J. Biol. Turk Biyol. Derg. 2020; 44:121–131

4. Islam MT, Sarkar C, El-Kersh DM, et al. Natural products and their derivatives against coronavirus: A review of the non-clinical and pre-clinical data. Phytother. Res. 2020; ptr.6700

5. Chen H, Du Q. Potential Natural Compounds for Preventing SARS-CoV-2 (2019-nCoV) Infection. 2020;

6. Moorthy V, Henao Restrepo AM, Preziosi M-P, et al. Data sharing for novel coronavirus (COVID-19). Bull. World Health Organ. 2020; 98:150–150

7. Zhang K. Is traditional Chinese medicine useful in the treatment of COVID-19? Am. J. Emerg. Med. 2020;

8. Li Y, Liu X, Guo L, et al. Traditional Chinese herbal medicine for treating novel coronavirus (COVID-19) pneumonia: protocol for a systematic review and meta-analysis. Syst. Rev. 2020; 9:75

9. Liu C. Pay attention to situation of SARS-CoV-2 and TCM advantages in treatment of novel coronavirus infection. Chin. Herb. Med. 2020; 12:97–103

10. Yang Y, Islam MS, Wang J, et al. Traditional Chinese Medicine in the Treatment of Patients Infected with 2019-New Coronavirus (SARS-CoV-2): A Review and Perspective. Int. J. Biol. Sci. 2020; 16:1708–1717

11. Dong L, Hu S, Gao J. Discovering drugs to treat coronavirus disease 2019 (COVID-19). Drug Discov. Ther. 2020; 14:58–60

12. Alam A, Mohd F siddiqui, Nikhat I, et al. COVID-19: Current Knowledge, Disease Potential, Prevention and Clinical Advances. Turk. J. Biol. 2020;

13. Tambunan. n Silico Analysis of Envelope Dengue Virus-2 and Envelope Dengue Virus-3 Protein as the Backbone of Dengue Virus Tetravalent Vaccine by Using Homology Modeling Method. OnLine J. Biol. Sci. 2009; 9:6–16

14. López JA, Weilenman C, Audran R, et al. A synthetic malaria vaccine elicits a potent CD8(+) and CD4(+) T lymphocyte immune response in humans. Implications for vaccination strategies. Eur. J. Immunol. 2001; 31:1989–1998

15. Shahsavandi S, Ebrahimi MM, Sadeghi K, et al. Design of a heterosubtypic epitope-based peptide vaccine fused with hemokinin-1 against influenza viruses. Virol. Sin. 2015; 30:200–207

16. Bourdette DN, Edmonds E, Smith C, et al. A highly immunogenic trivalent T cell receptor peptide vaccine for multiple sclerosis. Mult. Scler. Houndmills Basingstoke Engl. 2005; 11:552–561

17. Knutson KL, Schiffman K, Disis ML. Immunization with a HER-2/neu helper peptide vaccine generates HER-2/neu CD8 T-cell immunity in cancer patients. J. Clin. Invest. 2001; 107:477–484

18. Petrovsky N, Brusic V. Computational immunology: The coming of age. Immunol. Cell Biol. 2002; 80:248–254

19. Brusic V, Bajic VB, Petrovsky N. Computational methods for prediction of T-cell epitopes--a framework for modelling, testing, and applications. Methods San Diego Calif 2004; 34:436–443

20. Nielsen M, Lundegaard C, Lund O, et al. The role of the proteasome in generating cytotoxic T-cell epitopes: insights obtained from improved predictions of proteasomal cleavage. Immunogenetics 2005; 57:33–41

21. Bhattacharya M, Sharma AR, Patra P, et al. Development of epitope-based peptide vaccine against novel coronavirus 2019 (SARS-COV-2): Immunoinformatics approach. J. Med. Virol. 2020; 92:618–631

22. Joshi A, Joshi BC, Mannan MA, et al. Epitope based vaccine prediction for SARS-COV-2 by deploying immuno-informatics approach. Inform. Med. Unlocked 2020; 19:100338

23. Abdelmageed MI, Abdelmoneim AH, Mustafa MI, et al. Design of a Multiepitope-Based Peptide Vaccine against the E Protein of Human COVID-19: An Immunoinformatics Approach. BioMed Res. Int. 2020; 2020:2683286

24. Shang J, Wan Y, Luo C, et al. Cell entry mechanisms of SARS-CoV-2. Proc. Natl. Acad. Sci. 2020; 117:11727–11734

25. Ou X, Liu Y, Lei X, et al. Characterization of spike glycoprotein of SARS-CoV-2 on virus entry and its immune cross-reactivity with SARS-CoV. Nat. Commun. 2020; 11:1620

26. Sievers F, Wilm A, Dineen D, et al. Fast, scalable generation of high-quality protein multiple sequence alignments using Clustal Omega. Mol. Syst. Biol. 2011; 7:539

27. Larsen MV, Lundegaard C, Lamberth K, et al. Large-scale validation of methods for cytotoxic T-lymphocyte epitope prediction. BMC Bioinformatics 2007; 8:424

28. Lundegaard C, Lund O, Nielsen M. Accurate approximation method for prediction of class I MHC affinities for peptides of length 8, 10 and 11 using prediction tools trained on 9mers. Bioinforma. Oxf. Engl. 2008; 24:1397–1398

29. Buchan DWA, Jones DT. The PSIPRED Protein Analysis Workbench: 20 years on. Nucleic Acids Res. 2019; 47:W402–W407

30. Giguère S, Drouin A, Lacoste A, et al. MHC-NP: predicting peptides naturally processed by the MHC. J. Immunol. Methods 2013; 400-401:30–36

31. Wang P, Sidney J, Kim Y, et al. Peptide binding predictions for HLA DR, DP and DQ molecules. BMC Bioinformatics 2010; 11:568

32. Chou PY, Fasman GD. Prediction of the secondary structure of proteins from their amino acid sequence. Adv. Enzymol. Relat. Areas Mol. Biol. 1978; 47:45–148

33. Larsen J, Lund O, Nielsen M. [No title found]. Immunome Res. 2006; 2:2

34. Chou PY, Fasman GD. Empirical Predictions of Protein Conformation. Annu. Rev. Biochem. 1978; 47:251–276

35. Emini EA, Hughes JV, Perlow DS, et al. Induction of hepatitis A virus-neutralizing antibody by a virus-specific synthetic peptide. J. Virol. 1985; 55:836–839

36. Karplus PA, Schulz GE. Prediction of chain flexibility in proteins: A tool for the selection of peptide antigens. Naturwissenschaften 1985; 72:212–213

37. Kolaskar AS, Tongaonkar PC. A semi-empirical method for prediction of antigenic determinants on protein antigens. FEBS Lett. 1990; 276:172–174

38. Parker JMR, Guo D, Hodges RS. New hydrophilicity scale derived from high-performance liquid chromatography peptide retention data: correlation of predicted surface residues with antigenicity and x-ray-derived accessible sites. Biochemistry 1986; 25:5425–5432

39. Bui H-H, Sidney J, Li W, et al. Development of an epitope conservancy analysis tool to facilitate the design of epitope-based diagnostics and vaccines. BMC Bioinformatics 2007; 8:361

40. Calis JJA, Maybeno M, Greenbaum JA, et al. Properties of MHC class I presented peptides that enhance immunogenicity. PLoS Comput. Biol. 2013; 9:e1003266

41. Thévenet P, Shen Y, Maupetit J, et al. PEP-FOLD: an updated de novo structure prediction server for both linear and disulfide bonded cyclic peptides. Nucleic Acids Res. 2012; 40:W288–293

42. Ho BK, Brasseur R. The Ramachandran plots of glycine and pre-proline. BMC Struct. Biol. 2005; 5:14

43. Smith KJ, Reid SW, Harlos K, et al. Bound water structure and polymorphic amino acids act together to allow the binding of different peptides to MHC class I HLA-B53. Immunity 1996; 4:215–228

44. Rist MJ, Theodossis A, Croft NP, et al. HLA Peptide Length Preferences Control CD8 + T Cell Responses. J. Immunol. 2013; 191:561–571

45. Murthy VL, Stern LJ. The class II MHC protein HLA-DR1 in complex with an endogenous peptide: implications for the structural basis of the specificity of peptide binding. Struct. Lond. Engl. 1993 1997; 5:1385–1396

46. Berman HM. The Protein Data Bank. Nucleic Acids Res. 2000; 28:235–242

47. Bui H-H, Sidney J, Dinh K, et al. Predicting population coverage of T-cell epitope-based diagnostics and vaccines. BMC Bioinformatics 2006; 7:153

48. Fleri W, Vaughan K, Salimi N, et al. The Immune Epitope Database: How Data Are Entered and Retrieved. J. Immunol. Res. 2017; 2017:5974574

49. Dhanda SK, Mahajan S, Paul S, et al. IEDB-AR: immune epitope database-analysis resource in 2019. Nucleic Acids Res. 2019; 47:W502–W506

50. Grifoni A, Sidney J, Zhang Y, et al. A Sequence Homology and Bioinformatic Approach Can Predict Candidate Targets for Immune Responses to SARS-CoV-2. Cell Host Microbe 2020; 27:671–680.e2

51. Ahmed SF, Quadeer AA, McKay MR. Preliminary Identification of Potential Vaccine Targets for the COVID-19 Coronavirus (SARS-CoV-2) Based on SARS-CoV Immunological Studies. Viruses 2020; 12:254

52. Saha S, Raghava GPS. AlgPred: prediction of allergenic proteins and mapping of IgE epitopes. Nucleic Acids Res. 2006; 34:W202–209

53. Gupta S, Kapoor P, Chaudhary K, et al. In silico approach for predicting toxicity of peptides and proteins. PloS One 2013; 8:e73957

54. Friesner RA, Murphy RB, Repasky MP, et al. Extra Precision Glide: Docking and Scoring Incorporating a Model of Hydrophobic Enclosure for Protein−Ligand Complexes. J. Med. Chem. 2006; 49:6177–6196

55. Friesner RA, Banks JL, Murphy RB, et al. Glide: A New Approach for Rapid, Accurate Docking and Scoring. 1. Method and Assessment of Docking Accuracy. J. Med. Chem. 2004; 47:1739–1749

56. Halgren TA, Murphy RB, Friesner RA, et al. Glide: A New Approach for Rapid, Accurate Docking and Scoring. 2. Enrichment Factors in Database Screening. J. Med. Chem. 2004; 47:1750–1759

57. Halgren TA. Identifying and Characterizing Binding Sites and Assessing Druggability. J. Chem. Inf. Model. 2009; 49:377–389

58. Cohen AD, Shoenfeld Y. Vaccine-induced autoimmunity. J. Autoimmun. 1996; 9:699–703

59. Jain S, Baranwal M. Computational analysis in designing T cell epitopes enriched peptides of Ebola glycoprotein exhibiting strong binding interaction with HLA molecules. J. Theor. Biol. 2019; 465:34–44

60. Maslak PG, Dao T, Bernal Y, et al. Phase 2 trial of a multivalent WT1 peptide vaccine (galinpepimut-S) in acute myeloid leukemia. Blood Adv. 2018; 2:224–234

61. Sundar R, Rha SY, Yamaue H, et al. A phase I/Ib study of OTSGC-A24 combined peptide vaccine in advanced gastric cancer. BMC Cancer 2018; 18:332

62. Melief CJM, van der Burg SH. Immunotherapy of established (pre)malignant disease by synthetic long peptide vaccines. Nat. Rev. Cancer 2008; 8:351–360

63. Chiou S-S, Fan Y-C, Crill WD, et al. Mutation analysis of the cross-reactive epitopes of Japanese encephalitis virus envelope glycoprotein. J. Gen. Virol. 2012; 93:1185–1192

64. Alam A, Ali S, Ahamad S, et al. From ZikV genome to vaccine: in silico approach for the epitope-based peptide vaccine against Zika virus envelope glycoprotein. Immunology 2016; 149:386–399

65. He Y, Zhou Y, Wu H, et al. Identification of immunodominant sites on the spike protein of severe acute respiratory syndrome (SARS) coronavirus: implication for developing SARS diagnostics and vaccines. J. Immunol. Baltim. Md 1950 2004; 173:4050–4057

66. Yang J, James E, Roti M, et al. Searching immunodominant epitopes prior to epidemic: HLA class II-restricted SARS-CoV spike protein epitopes in unexposed individuals. Int. Immunol. 2009; 21:63–71

67. ohn Sidney; Jason Botten; Benjamin Neuman; Michael Buchmeier; Alessandro Sette. HLA DRB1*01:01 binding capacity of selected SARS-derived peptides. IEDB_SUBMISSION 2006;

68. Wang B, Chen H, Jiang X, et al. Identification of an HLA-A*0201–restricted CD8+ T-cell epitope SSp-1 of SARS-CoV spike protein. Blood 2004; 104:200–206

69. Guo J-P, Petric M, Campbell W, et al. SARS corona virus peptides recognized by antibodies in the sera of convalescent cases. Virology 2004; 324:251–256

