## Supplementary figures and images for "Design of an Epitope-Based Peptide Vaccine against the Severe Acute Respiratory Syndrome Coronavirus-2 (SARS-CoV-2): A Vaccine-informatics Approach"

### Suppli_Fig 1

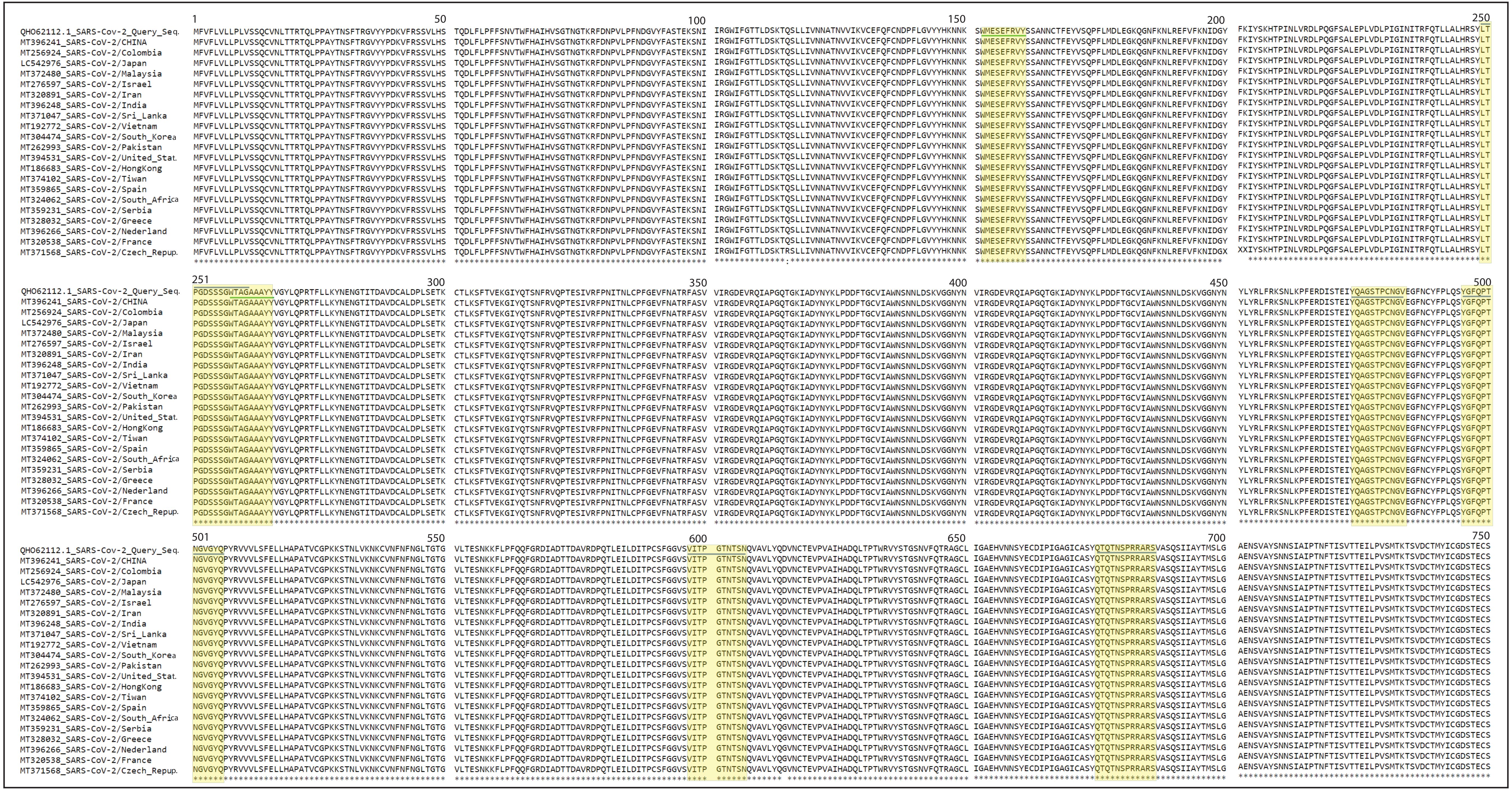

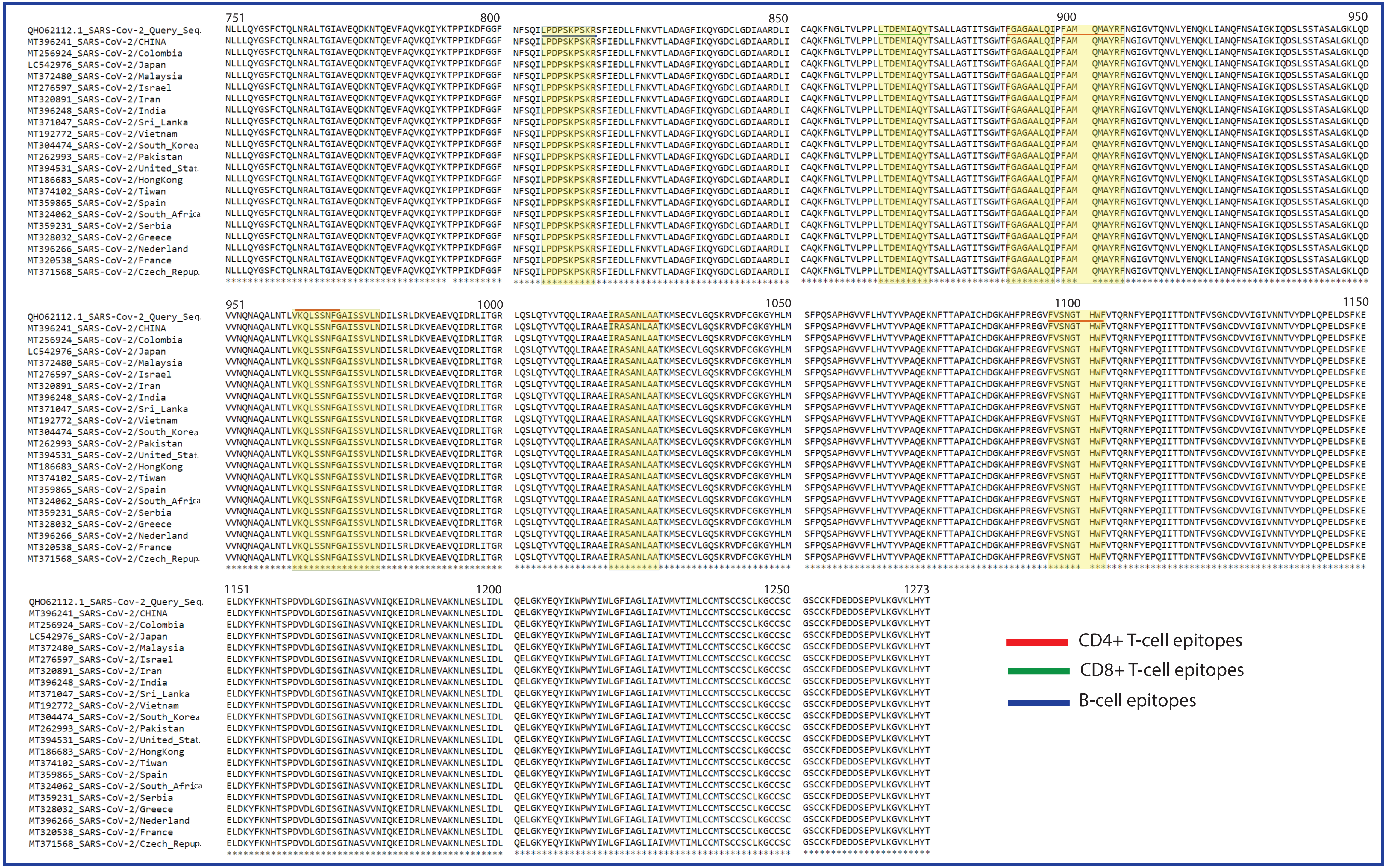
